# Nanopore-based analysis unravels the genetic landscape and phylogenetic relationships of human-infecting *Trichuris incognita* and *Trichuris trichiura* in Côte d’Ivoire, Uganda, Tanzania, and Laos

**DOI:** 10.1101/2024.07.31.605962

**Authors:** Nurudeen Rahman, Max A. Bär, Julian Dommann, Eveline Hürlimann, Jean T. Coulibaly, Said Ali, Somphou Sayasone, Prudence Beinamaryo, Jennifer Keiser, Pierre H.H. Schneeberger

## Abstract

Soil-transmitted helminthiases, particularly trichuriasis, affect over 500 million people, mostly in low- and middle-income countries. Traditional diagnostics fail to distinguish between *Trichuris* species, obscuring transmission patterns and treatment outcomes. Using nanopore-based full-length ITS2 rDNA sequencing, we analyzed 687 samples from Côte d’Ivoire, Laos, Tanzania, and Uganda, confirming the phylogenetic placement of *Trichuris trichiura* and the recently described *Trichuris incognita*. We identified two genetically distinct *Trichuris* species infecting humans, with divergent geographic patterns and presence in non-human primates, suggesting complex host-parasite dynamics. Within-country genetic variation indicated local adaptation and cryptic population structure. Importantly, we demonstrated that ITS2 fragment length is a robust, cost-effective diagnostic marker for differentiating *T. incognita* and *T. trichiura*, offering a practical alternative to sequencing for resource-limited settings. These findings expose the hidden complexity of *Trichuris* infections and highlight the urgent need to update diagnostic and control strategies to account for overlooked species diversity in endemic regions.

## Background

Trichuriasis, an infection caused by the whipworm *Trichuris spp*. is of considerable public health significance and is considered a Neglected Tropical Disease (NTD) ^1^. This particular NTD has been estimated to disproportionately infect over 465 million people worldwide, mainly prevalent among children living in impoverished communities with limited hygiene or sanitation facilities, leading to a significant burden of 0.64 million disability-adjusted life years (DALYs) lost annually ^2^. However, several other mammals such as non-human primates (NHP), rodents, ruminants, and marsupials to name a few encompass the more than 20 specific taxonomic groups of hosts associated specifically with the genus *Trichuris,* meaning they are multi-host parasites ^3,4^.

Previous research approaches using traditional diagnostic methods like Kato-Katz have always regarded *Trichuris sp*. found in humans and NHP as *Trichiura trichiura,* except in host-specific cases like the NHP Colobus monkeys (*Trichuris colobae*) ^5^, where there is a partial understanding of the *Trichuris* multi-host complexity. Morphologically, no reliable distinctions currently exist between the eggs or adult worms of *Trichuris* species infecting humans and non-human primates (NHP), making molecular methods essential for advancing our understanding of the genus *Trichuris* in humans and other hosts ^6–10^. Recent efforts employed mostly sequences from the internal transcribed spacer (ITS) regions 1 and 2 (ribosomal DNA), mitochondrial markers and β tubulin gene to delineate this cryptic difference, hence resolving the *Trichuris* complex or multi-host specificity ^5,8,9,11–15^. These sequences were used to prove the existence of two separate clusters of *Trichuris* in humans, with the consensus being that they both were *T. trichiura* ^8,9,11–16^. However, a recent study in Côte d’Ivoire revealed a novel *Trichuris* species, named *Trichuris incognita*, which forms a monophyletic clade genetically closer to *T. suis* than *T. trichiura* ^10,17^. ITS2 sequences corresponding to this species have been identified in earlier studies involving human patients from Uganda and Cameroon, as well as several non-human primates ^14^. However, in the absence of formal species classification, these sequences were previously labeled as *T. trichiura* or *Trichuris* sp. due to the absence of formal species classification. The ITS2 marker has also been shown to reliably distinguish between *T. suis* isolated from pigs, *T. incognita* isolated from humans, and *T. trichiura* isolated from humans and non-human primates ^10,17^.

In this study, we investigated the genetic diversity and phylogenetic placement of the human-infecting whipworms *T. incognita* and *T. trichiura* in four low- and middle-income countries (LMICs) across the globe. We analyzed the complete ITS2 region (∼500–600 bp) from whipworms found in stool samples collected from 687 patients participating in clinical studies conducted in Côte d’Ivoire, Tanzania, Uganda, and Laos. These samples were originally collected as part of randomized controlled trials assessing the efficacy of anthelmintic treatments ^18,19^. Additionally, we integrated the data generated from our analyses with publicly available ITS2 sequences from the genus *Trichuris*, encompassing sequences derived from humans, non-human primates, pigs, livestock, and rodents from diverse geographic locations. We then evaluated the genetic structure and phylogenetic relationships both within our study populations and in a broader global context. Finally, we identified a PCR-based diagnostic marker based on ITS2 fragment length that reliably differentiates between the two known human-infecting species, *T. trichiura* and *T. incognita*.

## Results

### Description of the study population

The population studied consisted of patients from the Pujini and Dodo shehias on Pemba Island, Tanzania; the Kisoro and Kabale districts in Uganda; the Dabou and Jacqueville regions around the Lagunes district in Côte d’Ivoire; and the Nambak district and Luang Prabang province in Laos (Figure 1, (adapted from Keller et al. ^20^). This study included frozen faecal samples from participants aged 6 to 60 years in Côte d’Ivoire, Tanzania and Laos, as well as ethanol-preserved faecal samples from school-aged children (6-12 years) in Uganda.

**Figure 1:**
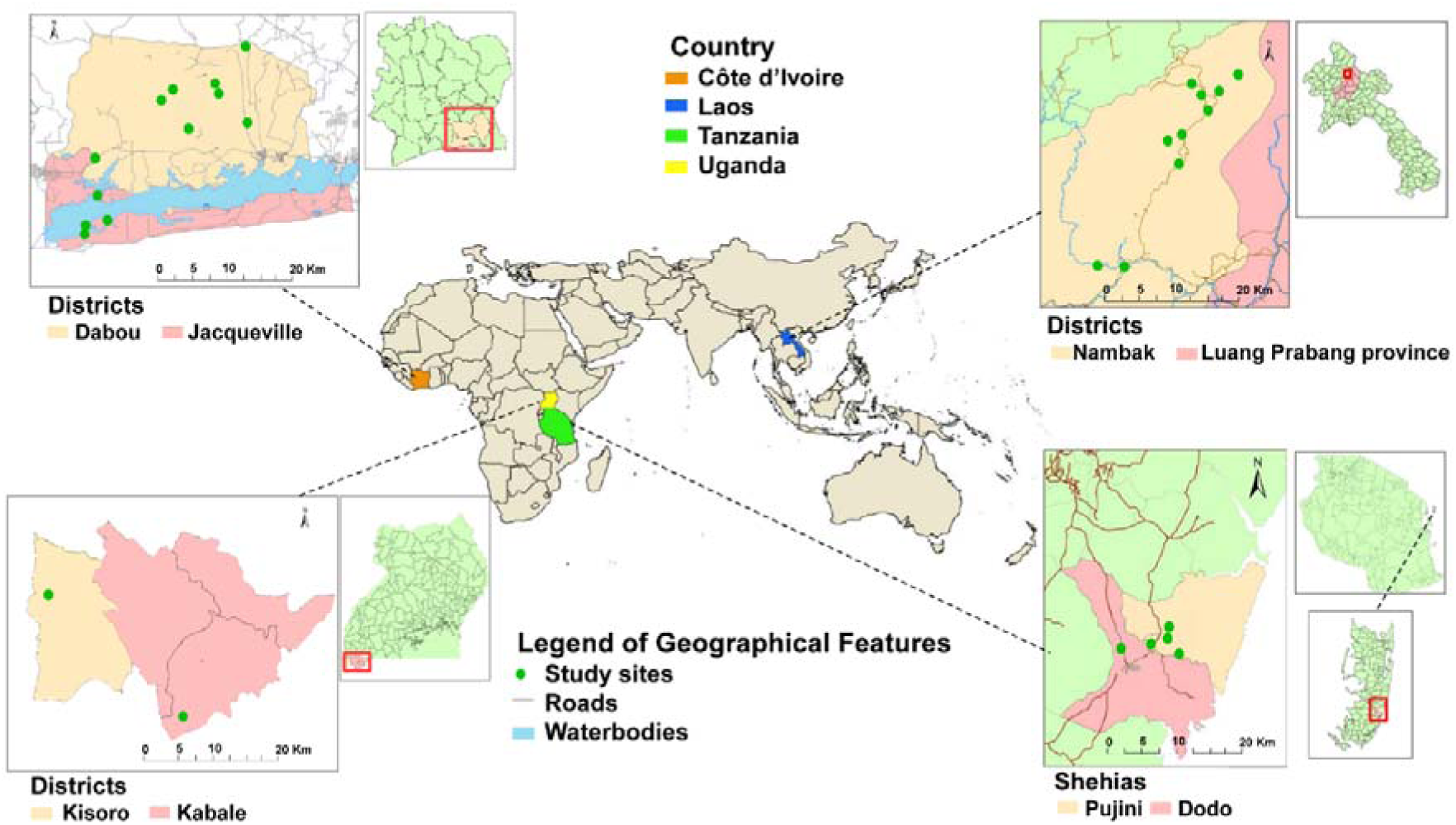
Sampling locations of the study population. Sampling locations of the study population. The map of the world highlights the four different study locations: the Lagunes region in Côte d’Ivoire, Pujini and Dodo Shehias on Pemba, Tanzania, Kisoro and Kabale district in Uganda, and the Nambak district and Luang Prabang province in Laos.

Stool samples used in this study were collected in the framework of a double-blind, placebo-controlled randomized trial conducted between September 2018 and June 2020 in Côte d’Ivoire, Tanzania, and Laos ^18,21,22^ as well as a parallel open-label randomized controlled superiority trial conducted between October to November 2023 in Uganda (registered as NCT06037876; clinicaltrials.gov/). Demographics and helminth infection characteristics of the participants included in this study are summarized in Table 1, Supplementary Data 1, and Supplementary Data 2.

**Table 1.** Description of the study population.

|  | Côte d'Ivoire<br>(n=186) | Laos<br>(n=173) | Tanzania<br>(n=188) | Uganda<br>(n=135) |
| --- | --- | --- | --- | --- |
| **Age in Years** |  |  |  |  |
| 6 -12 | 112 (60.2%) | 78 (45.1%) | 120 (63.8%) | 135 (100%) |
| 13 – 24 | 33 (17.7%) | 16 (9.2%) | 47 (25.0%) | 0 |
| ≥25 | 41 (22.1%) | 79 (45.7%) | 21 (11.2%) | 0 |
| **Sex** |  |  |  |  |
| Female | 94 (50.5%) | 91 (52.6%) | 107 (56.9%) | 78 (57.7%) |
| Male | 92 (49.5%) | 82 (47.4%) | 81 (43.1%) | 57 (42.2%) |
| **Infection intensity\*** |  |  |  |  |
| Light | 144 (77.4%) | 149 (86.1%) | 144 (76.6%) | 118 (87.4%) |
| Moderate | 39 (21.0%) | 23 (13.3%) | 41 (21.8%) | 17 (12.6%) |
| Heavy | 3 (1.6%) | 1(0.6%) | 3 (1.6%) | 0 |

\* Infection intensity was classified according to mean egg per gram (EPG) into light (1–999 EPG), moderate (1000–9999 EPG), and heavy (≥10 000 EPG). Five samples that only had country data (2 from Côte d'Ivoire, 2 from Tanzania, and 1 from Laos), but were missing additional metadata, were also added to the sequencing workflow and analysis, but were excluded from this table

Of the 687 participants, 54.3% (370/682) were female and 45.7% (312/682) were male across the four countries. Overall, the Lao participants were older (mean age 23.9 years [SD 16.7]) than the other three cohorts (mean age 17.3 years [SD 14.2] in Côte d’Ivoire, 13.9 years [SD 10.0] in Tanzania, and 9.4 years [SD 2.1] in Uganda). The infection intensities as diagnosed with Kato-Katz and sex distribution of the study participants depict a good balance between groups. Of note, at the start of the PCR-based investigation, we also included five additional samples lacking metadata apart from country of origin (2 from Côte d’Ivoire, 2 from Tanzania, and 1 from Laos), hence the total number of participants analyzed for the ITS2 region to 687.

### Molecular Analysis of ITS2 Nuclear Marker

Following DNA extraction, we successfully produced amplicons from 186, 188, 135, and 173 participants in Côte d’Ivoire, Tanzania and Uganda, and Laos, respectively. Using a Promethion sequencing platform, we generated a total of 30,794,777 (322-812,627 unprocessed reads per sample). Using a stringent DADA2-based pipeline (Supplementary Figure 1), we retained a total of 23,445,373 reads (109-585,322 high-quality reads per sample, Supplementary Figure 2a-c). Samples with more than 500 high-quality, post-DADA2 reads (657/687, 95.6%) were retained. The DADA2 error plots are shown in Supplementary Figure 3. Using this dataset, a total of 2828 ASVs were generated, which were further clustered into 215 unique ASV clusters with consensus sequences using a 98.5% identity threshold. This 98.5% threshold was defined based on ASV variation observed in our positive controls, which consisted of single-worm heads from *T. trichiura* and *T. incognita*. 8/215 consensus sequences were further removed as they formed singleton clusters and the remaining 207 ASV clusters (Supplementary Data 3) were then used for population genetics and phylogenetic inferences.

### Haplotype network analysis

For the population network analysis, the 207 ASVs were grouped into 170 haplotypes using default thresholds in DnaSP (Supplementary Data 4). Overall, 48/170 haplotypes were found in two or more countries, while 122 haplotypes were found in one setting only (66, 17, 25, and 14 haplotypes unique to Côte d’Ivoire, Tanzania, Uganda, and Laos, respectively). Prevalence of all the haplotypes is shown in Supplementary Data 4. Several haplotypes were present in a substantial number of samples, following country-specific patterns. For instance, Hap 1 was found in 7.0% of the samples from Laos and Tanzania, but 4.0% in the other settings. Hap 37 was the dominant haplotype in Laos (13%) but was only present in 1.0%-9.0% of samples in the other countries. The BLAST results of all haplotypes are summarized in Supplementary Data 5.

The statistical parsimony haplotype network analysis revealed a clear structure separating all the *Trichuris* haplotypes into two main clusters (Figure 2), suggesting the existence of at least two divergent evolutionary lineages circulating across these populations. An overrepresentation of haplotypes from Côte d’Ivoire was found to cluster together with our *T. incognita* single worm controls (N=4), with the most abundant haplotype being Hap 1 (5.4%). The second cluster, which contained *T. trichiura* single worm controls, was structured around Hap 37, Hap 38, and Hap 39. The significant difference between both clusters is highlighted by the number of nucleotide differences between the two least dissimilar sequences from both clusters (113 SNPs), thus representing a large portion of the expected amplicon sizes for both species (between ∼520-600bp). Interestingly, we also observed that one haplotype, Hap 155, which showed a significant amount of differences (N=42) with its closest related sequence from the *T. incognita* cluster, thus suggesting further stratification of the haplotype population. Overall, the haplotype network suggests that *T. trichiura*-like haplotypes are diversified into distinct subpopulations, while *T. incognita*-like haplotypes are more homogeneous and might result from more recently diverged populations.

**Figure 2:**
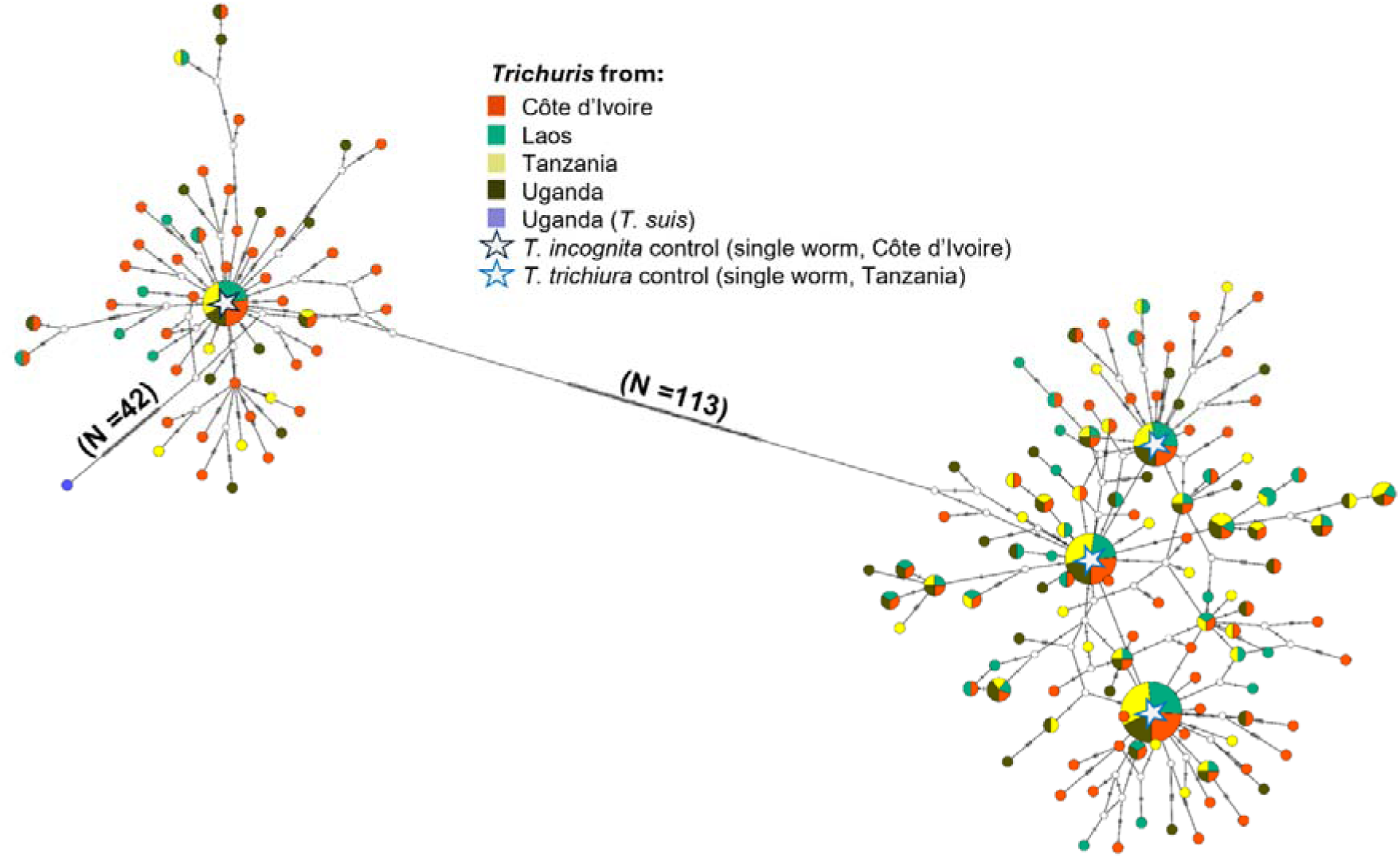
Underlying population genetic structure and network analysis among the four countries. Haplotype network inferred from ASVs of *Trichuris* species haplotype sequences from Côte d’Ivoire, Laos, Tanzania and Uganda. N = pairwise number of nucleotide differences between the main clusters; each hatch mark along the network branches corresponds to a nucleotide difference. Each circle represents a unique haplotype, and the circle’s size is proportional to the corresponding haplotype frequency. The colours of the pie charts represent the country of origin (Red for Côte d’Ivoire; green for Laos; yellow for Tanzania; dark green for Uganda). Star symbols denote reference control sequences: *Trichuris incognita* and *Trichuris trichiura* obtained from individual worms.

### Phylogenetic placement of sequence variants in the context of publicly available ITS2 sequences

The phylogenetic analysis was conducted using the 207 consensus sequences from ASV clusters generated in this study and a total of 97 ITS2 fragments obtained from GenBank. Firstly, to annotate each of our consensus sequences with a species label, we performed a clustering of all consensus sequences using a 90% threshold, which resulted in three main clusters, each containing the sequences extracted from the whole genome sequences of *T. trichiura*, *T. incognita*, and *T. suis*. Each consensus sequence was subsequently annotated with one of the 3 species, depending on which cluster they grouped in. To improve the clarity of the phylogenetic tree, we extracted the two most abundant consensus sequences corresponding to *T. trichiura* and *T. incognita* for samples with mixed infections, or the most abundant sequence in the case of a monoinfection, and used these sequences (N=1146 sequences) - along the public ITS2 references (N=97) – to generate the corresponding phylogenetic tree (Figure 3).

**Figure 3.**
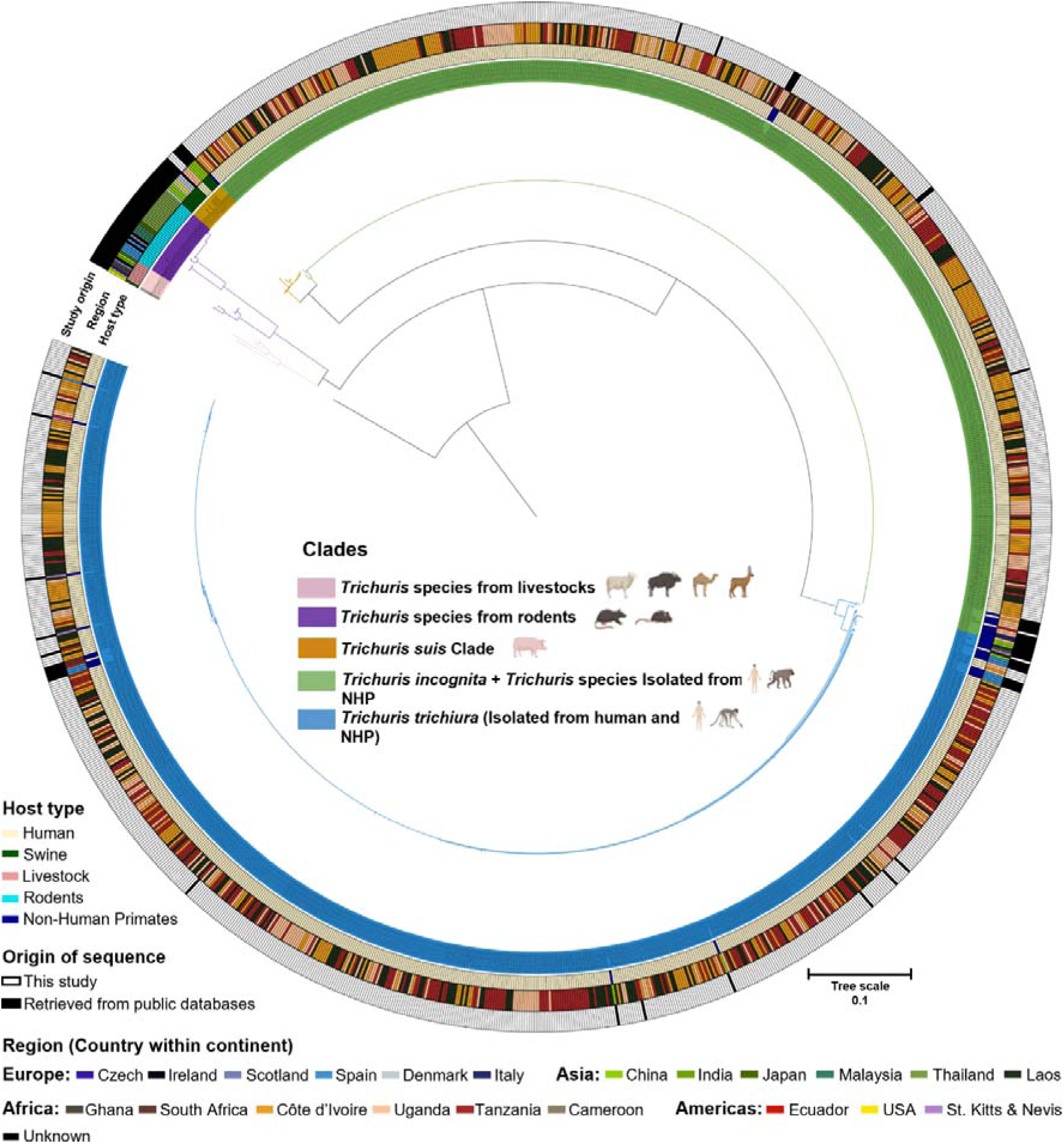
Phylogenetic analysis of ITS2 amplicons derived from *Trichuris-*positive stool samples of patients from Côte d’Ivoire, Laos, Tanzania, and Uganda. Maximum likelihood tree based on the ITS2 rDNA using Tamura-Nei with gamma distribution as the substitution model and *Trichinella spiralis* (GenBank accession: KC006432) as an outgroup. Colored branches correspond to major clades A to E, which broadly align with WGS-derived sequences from *T. trichiura*, *T. incognita*, *T. suis*, *T. muris*, and livestock-derived reference sequences. The external rings (from inner to outer) are colored according to metadata, including host type, country of origin, and whether the sequences were generated in this study or obtained from a public database.

We identified five clades, each corresponding to different but taxonomically related host species and broadly matching with published whole genome sequences (WGS) of *Trichuris* species. Clade A consisted of published ITS2 sequences of *Trichuris* species of Indian bison, camel, deer and sheep, and was the only clade which did not contain any well-characterized WGS of a specific *Trichuris* species. Clade B consisted of published sequences of *Trichuris* species of arvicolids and murid rodents, and included the sequence of an ITS2 region from a previously published WGS of *T. muris*. Clade C consisted of published sequences of *Trichuris* species of pigs, which correctly matched the WGS of *T. suis* ^23^. Interestingly, one of the sequences generated in this study, which was found in 3 samples from Uganda, clustered together with the WGS of *T. suis* ^23^ along with other *T. suis* sequences from China, Denmark and the USA. Clade D contains ITS2 sequences found in 44.8% of samples included in this study (N = 510 samples, with mono-or mixed infections), as well as 14 publicly available sequences of human and NHP species clustering with the WGS sequence of the novel species of human-infecting *Trichuris*, named *T. incognita*^10^. This clade also contained the sequences from the single worm positive controls from Côte d’Ivoire which we included in this study, as well as other published partial sequences from humans in Cameroon, non-human primates (NHP) from Italy, Uganda, South Africa. 626 sequences from our study clustered together with 36 ITS2 reference sequences of human and NHP species, four single worm controls from *T. trichiura*, and one WGS-derived sequence from the human type-species, *T. trichiura*, into clade E. Clade E also contained published sequences from humans in Ecuador and Uganda, as well as sequences from captive and non-captive NHP from Uganda, China, South Africa, Italy, Spain, and St. Kitts & Nevis.

Analysis of genetic diversity indices across *T. trichiura* and *T. incognita* populations in the four countries revealed considerable variation between species and sampling locations (Table 2).

**Table 2:** Genetic diversity estimations of *Trichuris* population per region.

| Country | Species | ASV clusters (N) | Nucleotide diversity ($\pi$) | Tajima's D | Fu and Li's D | Fu and Li's F |
| --- | --- | --- | --- | --- | --- | --- |
| Côte d'Ivoire | *T. trichiura* | 84 | 0.021 | -2.784** | -6.633** | -5.994** |
|  | *T. incognita* | 40 | 0.017 | -2.567*** | -4.278** | -4.252** |
| Laos | *T. trichiura* | 57 | 0.012 | -2.525** | -4.609** | -4.496** |
|  | *T. incognita* | 12 | 0.027 | -1.897* | -2.111* | -2.083* |
| Tanzania | *T. trichiura* | 58 | 0.011 | -2.541 | -4.939 | -4.748 |
|  | *T. incognita* | 11 | 0.012 | -2.111** | -2.570** | -2.463** |
| Uganda | *T. trichiura* | 55 | 0.015 | -2.725*** | -5.240** | -5.025** |
|  | *T. incognita* | 14 | 0.020 | -1.952* | -2.322 | -2.314* |

\* $p < 0.05$ ; \*\* $p < 0.01$ ; \*\*\* $p < 0.001$ (statistical significance levels)

Nucleotide diversity (π) varied notably across populations, with *T. incognita* from Laos (π = 0.027) and Uganda (π = 0.020) displaying the highest values, suggesting well-established populations with divergent haplotypes. In contrast, *T. trichiura* from Tanzania (π = 0.011) and Laos (π = 0.012) showed relatively low nucleotide diversity, indicative of recent population expansions or limited long-term genetic divergence. The *T. trichiura* population from Côte d’Ivoire demonstrated high diversity (π = 0.021) combined with a large number of ASV clusters (84), reflecting possible lineage mixing or stable long-term persistence. Interestingly, *T. incognita* in Laos exhibited fewer ASV clusters (12) but comparatively high nucleotide diversity, highlighting the presence of distinct and divergent haplotypes. Neutrality tests showed significantly negative Tajima’s D, Fu and Li’s D, and F values for *T. incognita* in Côte d’Ivoire, Laos, and Tanzania (*p* < 0.05), suggesting recent expansion or purifying selection. *T. trichiura* also showed similar patterns, except in Tanzania (*p* > 0.05).

Pairwise population comparisons based on F_ST_ and exact tests of sample differentiation revealed a marked contrast between species (Figure 4a-b). For *T. incognita*, some comparisons yielded low or negative F_ST_ values (e.g., LA vs TA: –0.0340; UG vs LA: – 0.0053), indicating little to no population differentiation. However, modestly elevated values were observed in comparisons involving Côte d’Ivoire, specifically CI vs TA (F_ST_= 0.0330, *p* < 0.05) and CI vs UG (F_ST_ = 0.0099) suggesting some degree of localized genetic divergence. In contrast, *T. trichiura* showed consistently low F_ST_ values (range: –0.0083 to – 0.0040), all of which were statistically non-significant (p > 0.05) apart from CI vs TA (F_ST_ = - 0.0040, *p* < 0.05), consistent with high gene flow across populations. These results suggest that *T. trichiura* populations remain relatively undifferentiated across countries, while *T. incognita* may exhibit more structured diversity, potentially reflecting recent expansion or geographic barriers to gene flow.

**Figure 4.**
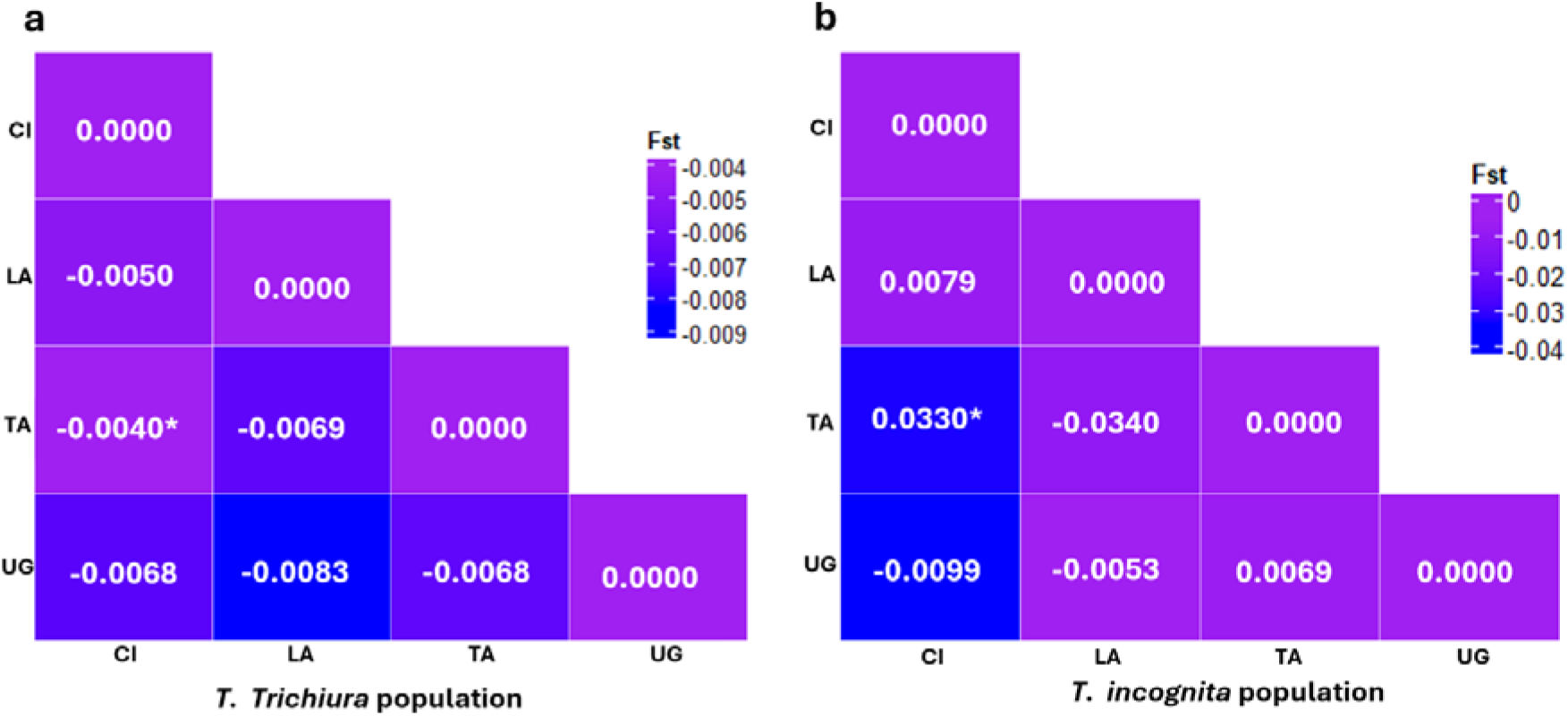
Pairwise F_ST_ comparisons among *Trichuris* populations from different regions. **a**. Visual representation of the matrix of pairwise F_ST_ comparisons among *Trichuris trichiura* populations across countries, indicating genetic differentiation. **b**. Visual representation of the matrix of pairwise F_ST_ comparisons among *Trichuris incognita* populations across countries, indicating genetic differentiation; CI = Côte d’Ivoire; LA = Laos; TA = Tanzania; UG = Uganda; (\**p* < 0.05)

### Comparative Sequence Analysis

We then conducted a comparative analysis across the clades and subclades previously defined for *Trichuris* spp. from humans, NHPs, rodents, livestock, and swine to assess intraspecific and interspecific similarities between *T. trichiura*, *T. incognita*, and *T. suis* (Figure 5a).

**Figure 5:**
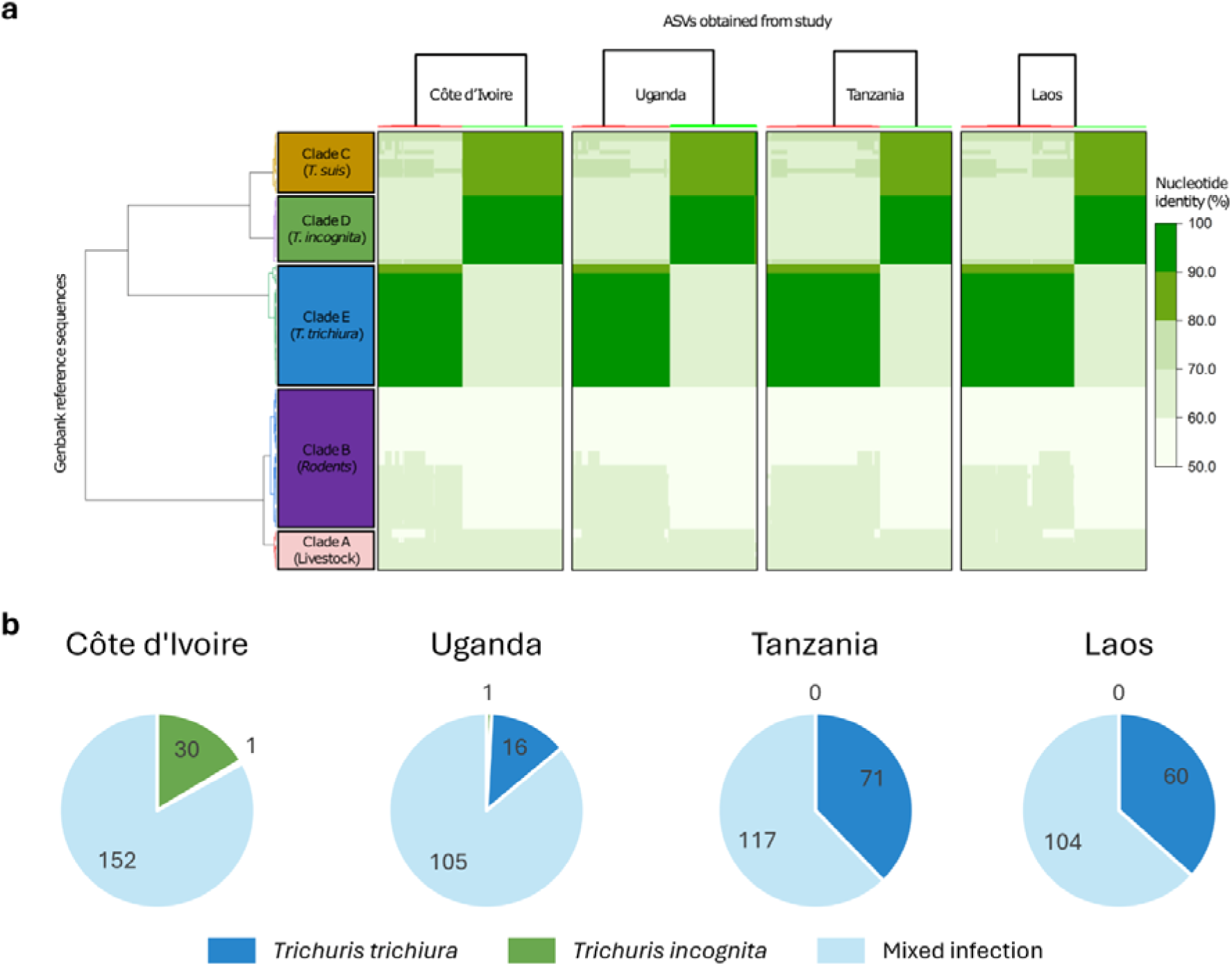
Comparative sequence analysis and proportion of *Trichuris* infection among the four countries. **a.** Pairwise comparison of sequence similarity between human samples (x-axis) from the four countries and reference sequences (y-axis). The reference sequences were clustered based on pairwise nucleotide identity values using the Ward clustering method. Sequences generated in this study were clustered separately by country using the same method. **b.** Prevalence of mono- and mixed infections comprising human-infecting *Trichuris trichiura* and *Trichuris incognita,* stratified by country.

By examining the pairwise genetic distance obtained across the four countries, it revealed a high similarity between the population of *Trichuris* across different clades. Investigation of the ASVs in the Clade D showed approximately 98%-100 % nucleotide sequence identity with the WGS-derived ITS2 sequence of *T. incognita*, observed in Côte d’Ivoire, as well as other published partial sequences from humans in Cameroon (accession number GQ301555), non-human primates (NHP) from Italy and Uganda, and more distantly with Chacma baboons in South Africa (accession number GQ301554) with 90% to 94% nucleotide sequence identity. Within Clade E (*T. Trichiura* lineage), we found a high similarity with populations of *T. trichiura* parasitizing humans from different geographical origins, as well as NHP with 96-100% nucleotide sequence identity. Furthermore, the *T. trichiura* clade was also more distant from clades C (*T. suis*) and D (*T. incognita*), with nucleotide identity values ranging from 64-69%. The single sequence from this study found to cluster together with *T. suis* in clade C shared nucleotide identities ranging from 97-99% identity with *T. suis* references, but only 82-88% identity and 70-72% identity with the *T. incognita* (clade D) and *T. trichiura* (clade E) clades, respectively. Analysis of the prevalence of human-infecting *Trichuris* species across the four countries revealed a high proportion of mixed infections, which accounted for the majority of cases in all settings, albeit with varying proportions (Figure 5b). Mixed infections accounted for 80.8% of cases in Côte d’Ivoire, 86.1% in Uganda, 63.4% in Tanzania, and 62.2% in Laos. Overall, *T. incognita* monoinfections were observed only in Côte d’Ivoire (16.4%) and Uganda (0.8%). *T. trichiura* monoinfections were detected in all countries, with the lowest prevalence in Côte d’Ivoire (0.5%), followed by Uganda (13.0%), Laos (36.6%), and the highest in Tanzania (37.8%). Based on sequence prevalence, we also generated relative abundance plots for samples with mixed infections, which followed a similar pattern to the overall prevalence: *T. incognita* relative abundance was highest in Côte d’Ivoire, followed by Uganda, and was lowest in Tanzania and Laos (Supplementary Figure 4).

### Fragment length as a robust diagnostic marker to differentiate human-infecting Trichuris species

The fragment lengths - in basepair (bp) - of the ITS2 regions corresponding to the clade structure (clades A to E) are summarized in Figure 6a. The length of the ITS2 regions ranged from 424.9±47.6bp for *T. muris*-related sequences, 443.7±7.9bp for *Trichuris* species found in livestock, 579.9±2.1bp for *T. suis*-related sequences, 595.9±1.5bp for *T. incognita*-related sequences, and 530.5±2.9bp for *T. trichiura-*related sequences. We observed significant differences in fragment length between all clades, except between *T. muris* and the livestock-related clade A. We next evaluated the predictive power of ITS2 region length for distinguishing between the two human-infecting species, *T. incognita* and *T. trichiura*. To this end, we applied a random forest classification model to our dataset, as illustrated in Figure 6b. As expected - given the complete lack of overlap in ITS2 fragment lengths between the two species - the model achieved perfect classification performance, with an area under the receiver operating characteristic curve (AUC) of 1.0, an accuracy of 100%, and a Cohen’s Kappa value of 1.0. The corresponding confusion matrix (Figure 6c) confirms a balanced distribution of data points across classes. Collectively, these findings demonstrate that ITS2 fragment length is a highly effective and reliable diagnostic marker for differentiating between *T. trichiura* and *T. incognita*.

**Figure 6:**
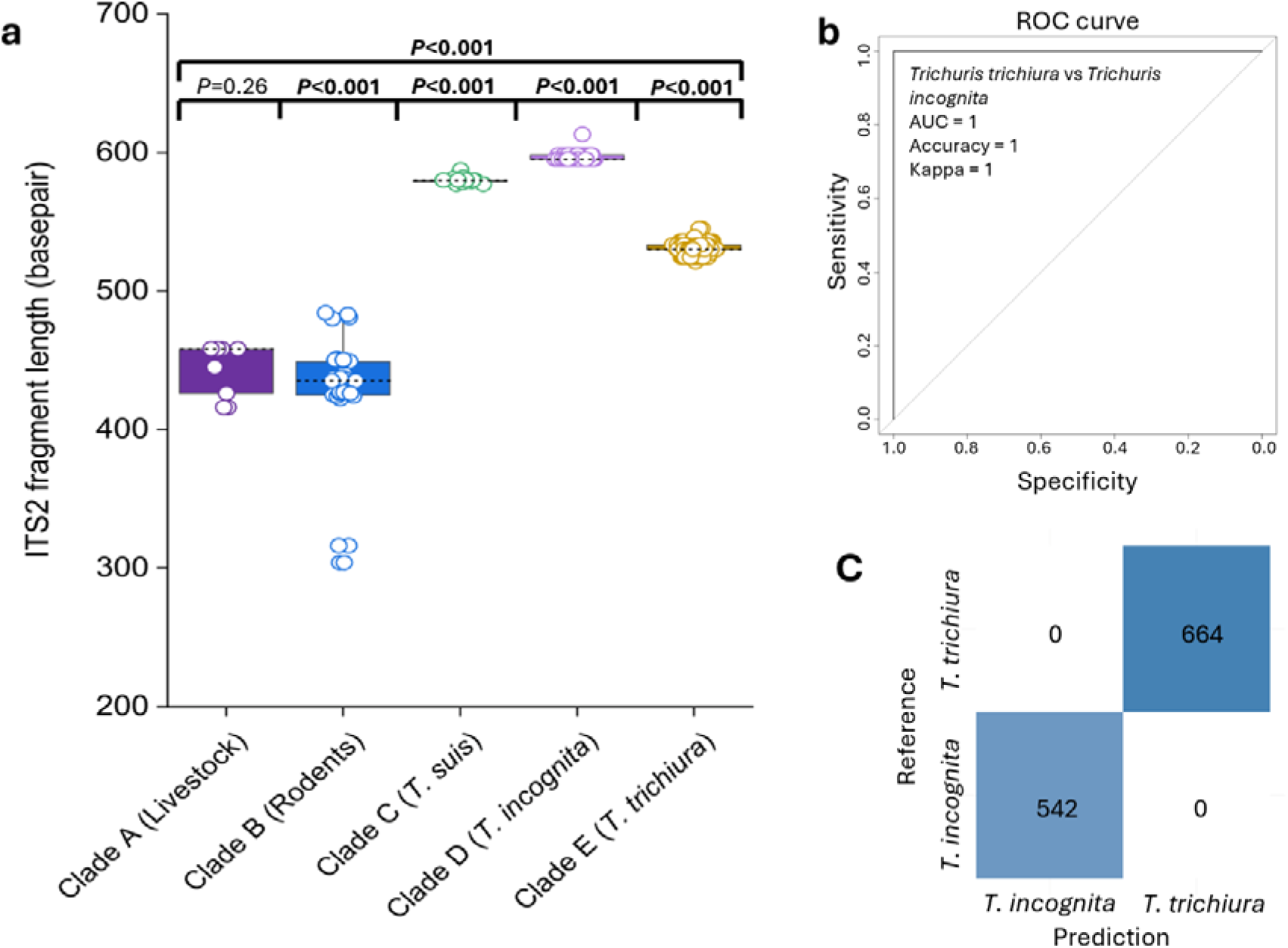
Fragment length could distinguish between different human-infecting *Trichuris* species. **a.** Comparison of the fragment length between the five clades. Box plots include the median line, and *p*-value estimated using pairwise Mann–Whitney tests. **b.** ROC curve showing the performance of a random forest model built using ITS2 fragment length, evaluated with a leave-one-out cross-validation (LOOCV) approach. **c.** Confusion matrix showing the class distribution of ITS2 fragment lengths from each *Trichuris* species. bp: base pair.

### Human-infecting Trichuris species and metadata associations

To explore potential associations between infection type and host characteristics, we focused on a subset of study participants aged 6–18 years (Supplementary Data 6). We investigated whether infections with *T. incognita* and *T. trichiura* were associated with differences in demographic or clinical metadata. Among infected individuals, females had a higher proportion of mixed infections in Tanzania (60.4%) and Uganda (55.5%). However, overall, there were no significant differences in the distribution of infection type (mixed vs. monoinfections with *T. incognita* or *T. trichiura*) by sex or country. Likewise, no significant differences were observed in infection intensity or egg counts between groups, suggesting that - within this age group - basic clinical and demographic parameters did not differ meaningfully between species (Figure 7c).

**Figure 7:**
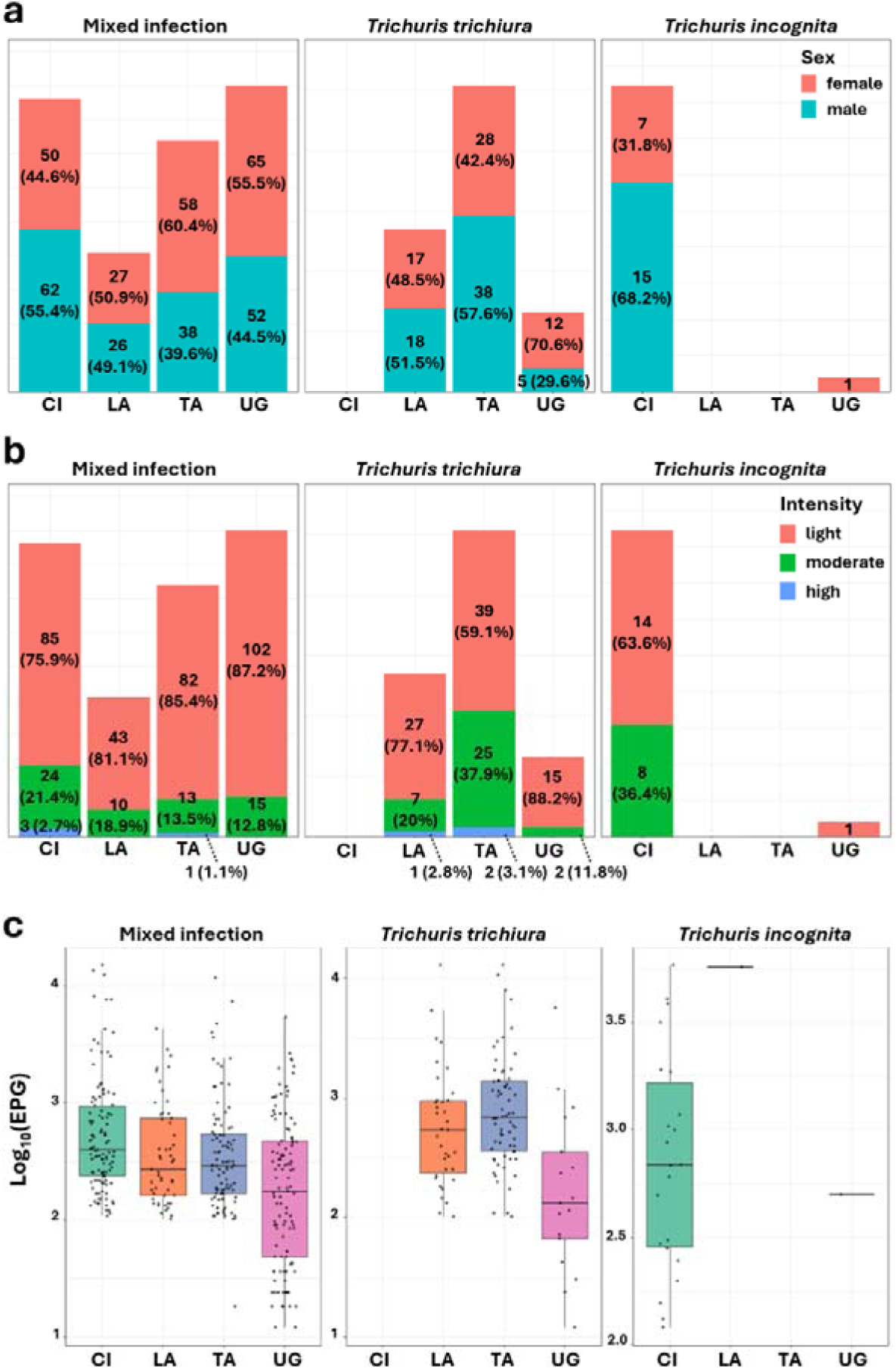
Demographic and infection characteristics of *Trichuris* spp. infections across four countries. **a.** Stacked bar plots showing the proportion of infected individuals by sex, with percentages indicating the proportion of male (turquoise) and female (red) participants within each infection type and country. **b.** Stacked bar plots showing the infection intensity distribution based on WHO criteria, with percentages representing the proportion of light (green), moderate (orange), and heavy (blue) infections among infected individuals in each category. **c.** Boxplots of log-transformed *Trichuris* egg counts per gram (EPG) of stool (right) for each infection type across countries. Each dot represents an individual.

## Discussion

Historically, the etiological agent of human trichuriasis has been identified as *Trichuris trichiura*, based on morphological characteristics^24^. However, morphology alone is often insufficient to reliably distinguish between whipworm species^24^. Recent molecular investigations, including whole-genome sequencing, have revealed a second *Trichuris* species infecting humans, *T. incognita*, which is morphologically indistinguishable from *T. trichiura* but displays significant genomic differences, including in the ITS regions^8,10,14,15,17,25,26^. Building on this discovery, we developed a nanopore-based amplicon sequencing strategy targeting the full ITS2 region to accurately identify *Trichuris* species in human samples collected through randomized controlled trials across diverse geographical regions.

Haplotype network analysis revealed a clear distinction between the two major *Trichuris* lineages infecting humans. Consistent with our earlier findings (Venkatesan et al.)^17^, we observed that most *T. incognita* sequences were derived from samples in Côte d’Ivoire, where the species was originally characterized. The nucleotide divergence between *T. incognita* and *T. trichiura* exceeded 30% in the ITS2 region, supporting their genetic distinctness. The haplotype network, centered on single-worm controls, suggested divergent evolutionary trajectories. While *T. incognita* haplotypes showed low regional structuring - indicative of recent spread or population homogeneity - *T. trichiura* showed broader diversity. A unique sequence diverging by 42 SNPs from *T. incognita* was also detected and phylogenetically grouped with *T. suis*, suggesting a rare zoonotic event or host-switching adaptation.

Our phylogenetic analysis of over 1,100 high-quality ITS2 sequences, combined with 97 public references, identified five well-supported clades consistent with host associations and whole genome-based *Trichuris* classifications. Human-derived sequences grouped into two genotypes: Clade D (*T. incognita*) and Clade E (*T. trichiura*). Clade D contained many sequences from Côte d’Ivoire samples, which were collected two years before the samples that led to the formal discovery of *T. incognita*^10^ in the same area, indicating that this species has likely been circulating undetected for some time. Notably, this is the first report of *T. incognita* in Tanzania, Laos, and Uganda, demonstrating a broader geographic distribution and significant diagnostic blind spots in microscopy-based detection.

Clade D also formed a monophyletic lineage with Clade C (*T. suis*), showing 82–88% nucleotide similarity. Although *T. incognita* and *T. suis* share morphological and ecological traits, their genetic divergence suggests parallel evolution from a distant common ancestor. *T. incognita* appears to be a multi-host lineage, as evidenced by its presence in 13 primate species across several countries and a human case from Cameroon^14^. ITS2 sequences across these hosts shared 98–100% identity, underscoring its zoonotic potential and possible existence in wildlife reservoirs - factors that complicate elimination efforts.

Clade E (*T. trichiura*) accounted for over 70% of sequences in this study and aligned with known human-derived sequences from Ecuador and Uganda^8,27^, as well as sequences from both captive and free-ranging non-human primates in Asia, Africa, and Europe^9,11,12,14,28^. These results reinforce previous findings that whipworms in primates form part of a complex *Trichuris* species network. Interestingly, three Ugandan human samples presented with sequences that clustered within the *T. suis* clade (Clade C), showing 97–99% identity to pig-derived sequences from China, Denmark, and the USA^13,23^. This points to a probable zoonotic transmission event or localized host switching, possibly influenced by gut microbiota^29^ - echoing observations by Nissen et al.^13^ in human–pig cohabitation zones. Although limited in number, these cases highlight the need to examine cross-species transmission in regions of intense human–livestock contact.

Our genetic diversity analyses revealed high sequence diversity and variable nucleotide diversity across species and locations. These patterns suggest the circulation of multiple lineages within both *T. trichiura* and *T. incognita*, potentially due to admixture, persistence, or secondary contact. Consistent with Venkatesan et al.^17^, we observed especially high diversity in Côte d’Ivoire. Slight discrepancies between our findings and theirs likely stem from our species-level sequence stratification, which reduces diversity signal due to the presence of multiple species. Overall, the data point to distinct evolutionary histories: *T. trichiura* exhibits broader diversity and moderate divergence, while *T. incognita* shows more localized, heterogeneous patterns potentially shaped by ecological or host-specific factors.

Placing these patterns into an evolutionary and public health context, our findings raise critical concerns. The frequent mixed infections, divergent population structures, and emergence of *T. incognita* in areas with suboptimal ivermectin-albendazole efficacy highlight the need for urgent reevaluation of species-specific transmission and treatment dynamics. With drug resistance already documented in veterinary whipworms^30–32^, emerging evidence of external variables such as the gut microbiota influencing treatment efficacy^33^, and signs of declining efficacy in human treatments^18^, the window to act preemptively is narrow.

To address this, deeper investigations are needed into strain-level variation, resistance markers, and ecological drivers of persistence, especially in regions where humans, non-human primates, and to some extent, pigs, share habitats. While high-throughput sequencing offers powerful resolution, it remains costly and impractical for routine diagnostics in low-resource settings^34,35^. To bridge this gap, we developed a PCR-based diagnostic marker based on ITS2 fragment length, which reliably distinguishes *T. trichiura* and *T. incognita*. This marker offers a cost-effective, scalable alternative to sequencing and can be easily implemented using conventional PCR or adapted for field-friendly platforms such as LAMP. Although our nanopore-based approach supports high-throughput analysis, length-based assays remain the most accessible solution for routine deployment.

Finally, clarifying the distribution, population structure, host range, and zoonotic potential of *Trichuris* species is essential for developing effective control and surveillance programs. These efforts are especially urgent in light of the WHO’s 2030 goal to eliminate trichuriasis as a public health problem^36^. Addressing this challenge will require coordinated One Health strategies that integrate human, animal, and environmental health perspectives into parasitic disease research and control.

### Limitations

Although our study included samples from four endemic regions, the geographic scope remains limited and may not fully capture the global diversity of the newly identified human-infecting *Trichuris* species. We also did not assess temporal factors, such as seasonal variation or long-term epidemiological trends. Environmental influences on the distribution and genetic structure of *Trichuris* populations were likewise not incorporated. To address these limitations, future studies should expand geographic and temporal sampling and integrate both nuclear and mitochondrial markers to enhance the resolution of population genetic analyses. Finally, we were unable to conduct association studies to explore potential links between *Trichuris* species, genetic structure, and treatment efficacy, as analyses from the primary clinical trials are still ongoing.

### Conclusion

This study presents the most comprehensive phylogenetic and diagnostic analysis of human-infecting *Trichuris* to date, revealing a wider distribution of *T. incognita* than previously recognized and demonstrating its zoonotic potential. By leveraging long-read ITS2 sequencing and introducing a robust, length-based diagnostic marker, we provide both foundational insight and practical tools to transform *Trichuris* surveillance and species-specific treatment strategies. These findings not only challenge existing diagnostic paradigms but also offer scalable solutions critical to achieving global trichuriasis elimination goals.

## Methods

### Study design and sample collection

The data and samples used in this study were derived from two studies. The first study was a double-blind, placebo-controlled randomized trial conducted between September 2018 and June 2020 in Côte d’Ivoire, Tanzania and Laos. The study examined the efficacy and safety of co-administered albendazole and ivermectin versus albendazole monotherapy against whipworm infections in children and adults with the rationale, procedures, and results of the study published previously ^18,21,22^.

Briefly, a fresh faecal sample was collected from community members aged between 6 and 60 years identified as infected with *T. trichiura* based on a prior stool examination and considered eligible for the trial with an infection intensity of at least 100 eggs per gram of stool. From every sample, about 1g of faeces was transferred into a 2ml screw cap cryotube using a UV sterilized plastic spatula and immediately frozen at −20°C, which were then shipped to Swiss TPH on dry ice at the end of the trial. Samples considered for this analysis represent baseline stool samples provided before any drug administration.

Additionally, data was also derived from a parallel open-label randomized controlled superiority trial conducted between October to November 2023 in two primary schools in Kabale and Kisoro districts, southwestern Uganda among individuals aged 6-12 years to investigate the superiority of co-administered ivermectin plus albendazole to albendazole monotherapy in terms of cure rates against *T. trichiura* infections (registered as NCT06037876; clinicaltrials.gov/). At baseline screening of the trial, two stool samples were collected at baseline and children found to be infected with *T. trichiura* based on quadruplicate Kato-Katz thick smear readings were enrolled into the study. A portion of the second baseline stool sample (1.5-2 g) from 135 eligible participants was preserved in 95% ethanol, shipped to Swiss TPH in Basel and subjected to amplicon sequencing for characterization of *Trichuris* species. Faecal egg count and epidemiological data are available in (Supplementary Table 3).

### Stool an worm DNA isolation and sample processing

Two separate extraction approaches were used in the isolation of DNA from stool received from the multi-country study and the study from Uganda. For the multi-country study, a total of 687 (188, 190, and 174 respectively from Côte d’Ivoire, Tanzania and Laos) frozen 100 mg of stool samples from the baseline were extracted using a QiAmp PowerFecal DNA extraction kit (Qiagen, Hilden, Germany) according to the manufacturer’s protocol with slight modification. For the faecal samples collected in Southwestern Uganda, 150 - 250mg of samples preserved in Ethanol was used in the extraction of the DNA. Before extraction with the QiAmp PowerFecal DNA extraction kit, a few modification steps were taken to mechanically rupture the *Trichuris* egg shells which include the removal of ethanol from the stool, washing with phosphate buffer saline (PBS), as well as a freeze-thaw-boiling step to enhance egg shell rupture and minimize inhibition ^17,37^ (see Supplementary Appendix 1 for more details of the modified ethanol-preserved sample extraction protocol).

DNA from adult worms received from two studies were prepared as follows: (i) for *T. incognita* samples collected from an expulsion study investigating the genomic characterization and discovery of *T. incognita* in Côte d’Ivoire, as previously described ^10^. DNA was extracted from anterior part of the worms using a DNeasy Blood & Tissue Kit (Qiagen, Cat: 69504) according to the manufacturer’s protocol; (ii) for the *T. trichiura* samples. The worms were collected as part of the Starworms project) in Tanzania which is Bill and Melinda Gates foundation funded (OPP1120972, PI is Bruno Levecke). DNA was also extracted from anterior part of the worms using a DNeasy Blood & Tissue Kit (Qiagen, Cat: 69504) according to the manufacturer’s protocol.

### PCR Amplification, Quantification and Purification

Primers used in the ITS2 rDNA reactions were *Trichuris*_ITS2_F 5’-ATGTCGACGCTACGCCTGTC-3’ and *Trichuris*_ITS2_R 5’-TAGCCTCGTCTGATCTGAGG-3’ ^17^ with a reverse primer containing an added custom barcodes that were adapted from the Illumina 12bp barcode sequences ^38,39^ (See Supplementary Data 7 for full primer sequences). For each run, PCR was carried out in 25 µl duplicate reactions to avoid bias. A 96-well plates were multiplexed with combined 2 μL of *Trichuris*_ITS2_F primers and eight custom barcoded reverse primers to amplify the ITS2-regions yielding unique barcodes of 592 bp ITS2 regions for each sample per plate column. 12.5 μL LongAmp® Hot Start Taq 2X Master Mix (cat no. M0533S, New England BioLabs, USA), 8.5 μL nuclease-free water and 2µL of source DNA were used in each amplification. Thermocycling conditions included the following: 94°C for 30 sec initially, 40 cycles of 94°C for 30 sec, 65°C for 15 sec, 65°C for 50 sec, followed by a final extension at 65°C for 10 min carried out on an Eppendorf Mastercycler Nexus Gradient Thermal Cycler (Eppendorf AG, Hamburg, Germany). Each pool of barcoded amplicons was quantified quantified directly using a dsDNA HS Assay kit on a Qubit 4 Fluorometer (Invitrogen, USA) according to the manufacturer’s protocol to ensure even throughput in the DNA pool using 2 µL of PCR reaction as input. 1% agarose and capillary gel electrophoresis (100V, 20min) were used to verify the amplification of the fragment at the expected band size. Afterwards, sample plates were stored overnight for a short period at 4°C overnight or at −20°C for long-term storage.

### Library preparation and sequencing

Nanopore sequencing libraries were prepared using the Native Barcoding Kit 24 v14 (SQK-NBD114.24, Oxford Nanopore Technologies, UK) according to the Native Barcoding Genomic DNA protocol with slight modifications aiming to normalize sample input, allowing for multiplexing as described in more detail elsewhere ^39^. Briefly, samples were pooled column-wise from a 96-well plate before the end-repair step, requiring 12 different outer barcodes for every 8 samples. Each column, representing 8 samples, was processed together with approximately 80 ng input corresponding to 200 fmol of the expected 0.59 kb fragments for the end-repair steps. End-prepped DNA was bead-cleaned with 0.7x ratio AMPure XP beads (Beckman Coulter) and resuspended in 15 μL nuclease-free water. For the “native barcoding ligation”, 7.50 μL end-prepped DNA was used instead of 0.75 μL, and barcoded samples were bead-purified using 0.6X ratio AMPure XP beads instead of 0.4X. For the “adapter ligation and clean-up” step, the pooled barcoded samples were bead-cleaned using 30 μL AMPure XP beads instead of 20 μL, utilizing a short-fragment buffer to preserve the ITS2 rDNA fragments. Afterwards, the final pooled library was quantified on the Qubit 4.0 (Invitrogen, USA) using the Qubit dsDNA HS Assay Kit (Invitrogen, USA), diluted in elution buffer from the kit to a total of ∼150 fmol and loaded onto the 10.4.1 p2 flow cells. Following this, sequencing was performed on the PromethION 2 Solo instrument (ONT) in four runs. Positive controls from individual worm heads of *Trichuris incognita* and *Trichuris trichiura* were also included in the runs.

### Read processing and sequencing analysis

All POD5 data resulting from the sequencing of the amplicons were already simplex basecalled through Dorado (v. 0.7.2; base-caller model dna_r10.4.1_e8.2_400bps_sup@v5.0.0), generating high-quality simplex reads with the – no-trim flag set. To ensure high-quality data for DADA2 ASV generation, demultiplexing by barcode using dorado (v0.9.6) for native barcodes with the--barcode-both-ends flag set, i.e., double-ended demultiplexing to reduce false positives. This was fed through our custom demultiplexing scripts that combine seqkit (v.2.6.1) ^40^ and custom Python scripts available at https://github.com/STPHxBioinformatics/HITS2. The script demultiplexes, quality filters, and length filters, as well as utilizes a minimum and maximum, trimmed-read cutoff of 520 bp and 750 bp, respectively, with zero mismatches allowed in primer recognition. Consequently, it trims the reads based on the inner PCR barcode (the reverse barcode in this case), ensuring the barcodes are found in the correct orientation. Afterwards, quality metrics were evaluated with NanoComp v1.12.0 ^41^ was then used for descriptive statistics on the runs. Demultiplexed ITS2 reads were then subjected to the R package ‘DADA2’ version 1.3.2 ^42^ pipeline to include further filtering to remove reads containing unresolved nucleotides (maxNLJ=LJ0) as well as reads exceeding the expected error number (maxEELJ=LJ3) and size range (520–700 bp). The dataset generated was then used as input to define the error rates and perform the removal of identical reads (derepFastq) while inferring the composition of the sequenced pool using the dereplicated sequence dataset as input (dadaFs), and removal of chimeric sequences. Following this, Amplicon sequence Variants (ASVs) generated through the DADA2 pipeline with ASVs below 5 reads across all samples were removed. After denoising with DADA2, all amplicon sequence variants (ASVs) were mapped against a reference database of *Trichuris* ITS2 sequences (Supplementary Data 7), using minimap2 (v2.24)^43^ to identify and remove off-target sequences. ASVs that did not align with at least 90% identity and 80% coverage to known *Trichuris* ITS2 references. Samples with less than 500 mapped reads were excluded from downstream analyses. To define an appropriate clustering threshold, we computed the pairwise average nucleotide identity (ANI) among ASVs derived from known positive controls (*T. trichiura* and *T. incognita* adult worm heads), yielding an average ANI of 98.7%. Based on this, we applied a 98.5% identity threshold using VSEARCH (v2.23.0)^44^ for intra-species clustering a data smoothing. Finally, to explore broader intraspecific and interspecific patterns, ASVs were clustered at 90% identity threshold to examine deeper divergence within and between species. An overview of the entire workflow is shown in Supplementary Figure 1. The Basic Local Alignment Search Tool (BLAST) available at GeneBank (https://www.ncbi.nlm.nih.gov/genbank/) and VSEARCH (v2.23.0)^44^ using our custom ITS2 reference sequence were used to verify correct species assignment and to fill in missing taxonomic data for unresolved ASVs based on identity.

### Genetic Variation and Phylogenetic Analysis

A curated database of reference ITS2 sequences was created by downloading all publicly available full ITS1-5.8S-ITS2 reference sequences of closely related species on NCBI using the search term “*Trichuris*”[porgn: txid36086] as well as from the WGS data of *Trichuris muris* ^27,45^, *Trichuris trichiura* ^27^, *Trichuris suis* ^23^ and an adult worm from CLJte d’Ivoire ^10^. Afterwards, seqkit ^40^ was used to extract only ITS2 sequences within the forward and reverse primers used in this study with the command seqkit-j 4 amplicon –m 2 -p primers.tab -r 1:-1 20240201_Trichuris.txid_36086.fasta--bed > 20240201_Trichuris_txid_36086.bed. The ASVs were then subjected to multiple sequence alignments with the newly curated reference sequences (Supplementary Table 8), as well as an outgroup species, *Trichinella spiralis* (Accession No: KC006432), using the MAFFT tool (Katoh et al., 2019). Phylogenetic relatedness was inferred using NJ in MEGA v11.0 ^46^ and the Maximum Likelihood method using RaxML v8.0 ^47^ with the “autoMRE bootstrapping”. Finally, the tree was visualized using the iTOL (interactive tree of life) software^48^.

### Population Genetic Structure

Alignment from good quality ITS2 ASVs was used for population genetic data and haplotype polymorphic analysis such as numbers of variable sites, number of haplotypes, nucleotide diversity (π), Tajima’s D, Fu and Li’s D and F’s between the *Trichuris* populations in the four countries were calculated using DnaSP v6.0 ^49^. Dnasp v6.0 ^49^ was also used to create haplotype files, including aligned variable nucleotides and information on the frequencies of each sequence. Statistical Parsimonious networks were then inferred and visualized using Population Analysis with Reticulate Trees, POPART v1.7 ^50^ with the connection limit set to 95% and gaps being treated as missing. We then calculated pairwise F_ST_ calculations in Arlequin v3.5.2 ^51^ using 1000 permutations. Statistical Parsimonious networks were inferred and visualized using POPART v1.7 ^50^ with the connection limit set to 95% and gaps being treated as missing. Furthermore, the number of nucleotide differences per sequence in each country and other *Trichuris* species incorporated in the phylogenetic analysis was analyzed using the Compute Pairwise Distance prompt based on the number of differences method of MEGA v11 ^46^ to assess the sequence similarity.

### Statistics analysis

To compare infection intensity and EPG values across sex and age groups across a subset of study participants aged 6–18 years. For the EPG, we applied a Kruskal–Wallis test (degrees of freedom dependent on the number of groups) followed by pairwise Wilcoxon rank sum exact tests for multiple comparisons (two-sided), using RStudio equipped with R version 4.4.2. These analyses were conducted on log-transformed EPG values, where appropriate, to reduce skewness. For categorical comparisons, such as sex across countries or age groups, Fisher’s exact tests were used, and *p*-values are reported in the results where relevant. The package randomForest v4.6–14^52^ was used to run random forest models using ITS2 fragment length, evaluated with a leave-one-out cross-validation (LOOCV) approach. The receiver operating characteristic (ROC) curves calculations were done using the pROC package v.1.16.2^53^. All graphs, besides the phylogenetic tree, were generated using the R version 4.4.2 and the OriginPro 2021 graphing software v9.8.0.200 (OriginLab Corporation, Northampton, MA, USA).

## Data Availability

The custom demultiplexing script and DADA2 analysis code used for this study are available at: https://github.com/STPHxBioinformatics/HITS2. The raw ONT sequence data (ITS2 rDNA amplicon sequencing) generated in this study is available in the NCBI Short Read Archive under accession (https://www.ncbi.nlm.nih.gov/sra; Bioproject no.PRJNA1283282).

## Acknowledgements

We want to acknowledge the participants and investigators of the IVM-ALB multi-country and FACE-IT efficacy trial study. We are also grateful to Bruno Levecke for providing the *Trichuris trichiura* worms, which served as essential positive controls to validate our molecular and bioinformatics methods. We are grateful for the support and access to the high-performance computing (HPC) cluster provided by the sciCORE scientific computing center at the University of Basel (http://scicore.unibas.ch/).

## Contributions

**NR**: study design, research design, project supervision, experimental work, statistical analyses, figure generation, writing of the first draft, and paper editing. **MB** and **JD**: provision of WGS from Côte d’Ivoire, and paper editing. **JC**, **EH**, **SA**, **PB**, and **SS**: study design, conducted fieldwork (sample collection, handling, parasitological work, and data curation), and paper editing. **JK**: study design, research design, project supervision, funding acquisition, and paper editing. **PHHS**: study design, research design, project supervision, funding acquisition, figure generation, statistical analysis, project supervision, and paper editing.

## Ethical declarations

The trial received approval from independent ethics committees in Côte d’Ivoire (reference numbers 088–18/MSHP/CNESVS-km and ECCI00918), Laos (reference number 093/NECHR), Tanzania (Zanzibar, reference number ZAMREC/0003/Feb/2018), Uganda (reference number HS3160ES/UNCST; Division of Vector-borne and neglected tropical diseases VCDR-2023-29) and the institutional research commission of the Swiss TPH and the ethics committee of Switzerland (EKNZ: Ethics Committee of North-Western and Central Switzerland; O_2023-00066; reference number BASEC Nr Req-2018-00494). The trial protocols have also been registered as NCT03527732 and NCT06037876 on ClinicalTrials.gov. Participants gave written informed consent (adults/parents) and assent (minors) to participation, and participants were allowed to withdraw from the study at any time point without further consequences.

## References

1. Bethony, J. et al. Soil-transmitted helminth infections: ascariasis, trichuriasis, and hookworm. The Lancet 367, 1521–1532 (2006).

2. Montresor A, Mwinzi P, Mupfasoni D, Garba A Reduction in DALYs lost due to soil-transmitted helminthiases and schistosomiasis from 2000 to 2019 is parallel to the increase in coverage of the global control programmes. PLoS Negl Trop Dis 16, e0010575 (2022).

3. Bundy, D. A. P. & Cooper, E. S. *Trichuris* and Trichuriasis in humans. Advances in Parasitology. 28, 107–173 (1989).

4. Callejón, R., Robles, M. D. R., Panei, C. J. & Cutillas, C. Molecular diversification of *Trichuris* spp. from *Sigmodontinae (Cricetidae)* rodents from Argentina based on mitochondrial DNA sequences. Parasitol Res 115, 2933–2945 (2016).

5. Cutillas, C., de Rojas, M., Zurita, A., Oliveros, R. & Callejón, R. *Tr*ichuris colobae n. sp. (Nematoda: Trichuridae), a new species of Trichuris from Colobus guereza kikuyensis. Parasitol Res 113, 2725–2732 (2014).

6. Oliveros, R., Cutillas, C., De Rojas, M. & Arias, P. Characterization of four species of *Trichuris* (Nematoda: Enoplida) by their second internal transcribed spacer ribosomal DNA sequence. Parasitol Res 86, 1008–1013 (2000).

7. Ooi, H.-K., Tenora, F., Itoh, K. & Kamiya, M. Comparative Study of *Trichuris trichiura* from non-human primates and from man, and their difference with *T. suis*. Journal of Veterinary Medical Science 55, 363–366 (1993).

8. Ghai, R. R. et al. Hidden population structure and cross-species transmission of whipworms (*Trichuris* sp.) in humans and non-human primates in Uganda. PLoS Negl Trop Dis 8, e3256 (2014).

9. Rivero, J., Cutillas, C. & Callejón, R. *Trichuris trichiura* (Linnaeus, 1771) From human and non-human Primates: morphology, biometry, host specificity, molecular characterization, and phylogeny. Front. Vet. Sci. 7, 626120 (2021).

10. Bär, M. A., et al. Genomic and morphological characterization of *Trichuris incognita*, a human-infecting *Trichuris* species. Preprint at 10.1101/2024.06.11.598441 (2025).

11. Cavallero, S. et al. Genetic heterogeneity and phylogeny of *Trichuris* spp. from captive non-human primates based on ribosomal DNA sequence data. *Infection*, Genetics and Evolution 34, 450–456 (2015).

12. Cavallero, S. et al. Nuclear and mitochondrial data on *Trichuris* from *Macaca fuscata* support evidence of host specificity. Life 11, 18 (2020).

13. Nissen, S. et al. Genetic analysis of *Trichuris suis* and *Trichuris trichiura* recovered from humans and pigs in a sympatric setting in Uganda. Veterinary Parasitology 188, 68–77 (2012).

14. Ravasi, D. F., O’Riain, M. J., Davids, F. & Illing, N. Phylogenetic evidence that two distinct *Trichuris* genotypes infect both humans and non-human primates. PLoS ONE 7, e44187 (2012).

15. Xie, Y. et al. Genetic characterisation and phylogenetic status of whipworms (*Trichuris* spp.) from captive non-human primates in China, determined by nuclear and mitochondrial sequencing. Parasites Vectors 11, 516 (2018).

16. Callejón, R., Halajian, A. & Cutillas, C. Description of a new species, Trichuris ursinus n. sp. (Nematoda: Trichuridae) from Papio ursinus Keer, 1792 from South Africa. Infection, Genetics and Evolution 51, 182–193 (2017).

17. Venkatesan A, Chen R, Bär M, Schneeberger P, Reimer B, Hürlimann E, et al. Trichuriasis in human patients from Côte d’Ivoire caused by novel species *Trichuris incognita* with low sensitivity to albendazole/ivermectin combination treatment. Emerg Infect Dis. 31:104–114. (2025).

18. Hürlimann, E. et al. Efficacy and safety of co-administered ivermectin and albendazole in school-aged children and adults infected with Trichuris trichiura in Côte d’Ivoire, Laos, and Pemba Island, Tanzania: a double-blind, parallel-group, phase 3, randomised controlled trial. The Lancet Infectious Diseases 22, 123–135 (2022).

19. Palmeirim MS, Hürlimann E, Beinamaryo P, Kyarisiima H, Nabatte B, Hattendorf J, et al. Efficacy and safety of albendazole alone *versus* albendazole in combination with ivermectin for the treatment of *Trichuris trichiura* infections: An open-label, randomized controlled superiority trial in south-western Uganda. PLoS Negl Trop Dis 18(11): e0012687.(2024).

20. Keller, L. et al. Insights gained from conducting a randomised controlled trial on ivermectin-albendazole against *Trichuris trichiura* in Côte d’Ivoire, Lao PDR and Pemba Island. Advances in Parasitology. 111: 253–276 (2021).

21. Keller, L. et al. Long-term outcomes of ivermectin-albendazole versus albendazole alone against soil-transmitted helminths: Results from randomized controlled trials in Lao PDR and Pemba Island, Tanzania. PLoS Negl Trop Dis 15, e0009561 (2021).

22. Patel, C. et al. Efficacy and safety of ivermectin and albendazole co-administration in school-aged children and adults infected with *Trichuris trichiura*: study protocol for a multi-country randomized controlled double-blind trial. BMC Infect Dis 19, 262 (2019).

23. Jex, A. R. et al. Genome and transcriptome of the porcine whipworm *Trichuris suis*. Nat Genet 46, 701–706 (2014).

24. Betson, M., Søe, M. J. & Nejsum, P. Human trichuriasis: Whipworm genetics, phylogeny, transmission and future research directions. Curr Trop Med Rep 2, 209–217 (2015).

25. Doyle, S.R., Søe, M.J., Nejsum, P. et al. Population genomics of ancient and modern *Trichuris trichiura*. Nat Commun 13, 3888 (2022.

26. Hawash, M. B. F. et al. Whipworms in humans and pigs: origins and demography. Parasites Vectors 9, 37 (2016).

27. Foth, B. J. et al. Whipworm genome and dual-species transcriptome analyses provide molecular insights into an intimate host-parasite interaction. Nat Genet 46, 693–700 (2014).

28. Liu, G.-H. et al. Mitochondrial and nuclear ribosomal DNA evidence supports the existence of a new *Trichuris* Species in the Endangered François’ Leaf-Monkey. PLoS ONE 8, e66249 (2013).

29. Stark, K. A. et al. Host microbiome determines host specificity of the human whipworm, Trichuris trichiura. Preprint at 10.1101/2025.05.09.653168 (2025).

30. Kelleher, A. C., Good, B., de Waal, T. & Keane, O. M. Anthelmintic resistance among gastrointestinal nematodes of cattle on dairy calf to beef farms in Ireland. Irish Veterinary Journal 73, 12 (2020).

31. Beleckė, A. et al. Anthelmintic resistance in small ruminants in the Nordic-Baltic region. Acta Veterinaria Scandinavica 63, 18 (2021).

32. Fairweather, I., Brennan, G. P., Hanna, R. E. B., Robinson, M. W. & Skuce, P. J. Drug resistance in liver flukes. International Journal for Parasitology: Drugs and Drug Resistance 12, 39–59 (2020).

33. Schneeberger, P.H.H., Gueuning, M., Welsche, S. et al. Different gut microbial communities correlate with efficacy of albendazole-ivermectin against soil-transmitted helminthiases. Nat Commun 13, 1063 (2022).

34. Su, C., Zhang, X. & Dubey, J. P. Genotyping of *Toxoplasma gondii* by multilocus PCR-RFLP markers: A high resolution and simple method for identification of parasites. International Journal for Parasitology 36, 841–848 (2006).

35. Francisco, C. J., Almeida, A., Castro, A. M. & Santos, M. J. Development of a PCR-RFLP marker to genetically distinguish *Prosorhynchus crucibulum* and *Prosorhynchus aculeatus*. Parasitology International 59, 40–43 (2010).

36. World Health Organization. Ending the Neglect to Attain the Sustainable Development Goals: A Road Map for Neglected Tropical Diseases 2021–2030. https://www.who.int/publications/i/item/9789240010352 (2021).

37. Ayana, M. et al. Comparison of four DNA extraction and three preservation protocols for the molecular detection and quantification of soil-transmitted helminths in stool. PLoS Negl Trop Dis 13, e0007778 (2019).

38. Caporaso, J. G. et al. Ultra-high-throughput microbial community analysis on the Illumina HiSeq and MiSeq platforms. The ISME Journal 6, 1621–1624 (2012).

39. Dommann, J. et al. A novel barcoded nanopore sequencing workflow of high-quality, full-length bacterial 16S amplicons for taxonomic annotation of bacterial isolates and complex microbial communities. mSystems 0, e00859–24 (2024).

40. Shen, W., Le, S., Li, Y. & Hu, F. SeqKit: A cross-platform and ultrafast toolkit for FASTA/Q file manipulation. PLoS ONE 11, e0163962 (2016).

41. Wouter De Coster, Rosa Rademakers, NanoPack2: population-scale evaluation of long-read sequencing data. Bioinformatics, 39, 5 (2023).

42. Callahan, B. J. et al. High-throughput amplicon sequencing of the full-length 16S rRNA gene with single-nucleotide resolution. Nucleic Acids Research 47, e103–e103 (2019).

43. Li, H. New strategies to improve minimap2 alignment accuracy. Bioinformatics 37, 4572– 4574 (2021).

44. Rognes, T., Flouri, T., Nichols, B., Quince, C. & Mahé, F. VSEARCH: a versatile open source tool for metagenomics. PeerJ 4, e2584 (2016).

45. Howe, K. L., Bolt, B. J., Shafie, M., Kersey, P. & Berriman, M. WormBase ParaSite − a comprehensive resource for helminth genomics. Molecular and Biochemical Parasitology 215, 2–10 (2017).

46. Tamura, K., Stecher, G. & Kumar, S. MEGA11: Molecular evolutionary genetics analysis version 11. Molecular Biology and Evolution 38, 3022–3027 (2021).

47. Stamatakis, A. RAxML version 8: a tool for phylogenetic analysis and post-analysis of large phylogenies. Bioinformatics 30, 1312–1313 (2014).

48. Letunic, I. & Bork, P. Interactive tree of life (iTOL) v6: recent updates to the phylogenetic tree display and annotation tool. Nucleic Acids Research 52, W78–W82 (2024).

49. Rozas, J. et al. DnaSP 6: DNA sequence polymorphism analysis of large data sets. Molecular Biology and Evolution 34, 3299–3302 (2017).

50. Leigh, J. W. & Bryant, D. popart: full-feature software for haplotype network construction. Methods in Ecology and Evolution 6, 1110–1116 (2015).

51. Excoffier, L. & Lischer, H. E. L. Arlequin suite ver 3.5: a new series of programs to perform population genetics analyses under linux and windows. Molecular Ecology Resources 10, 564–567 (2010).

52. Breiman, L. Random forests. Machine Learning 45, 5–32 (2001).

53. Robin, X. et al. pROC: an open-source package for R and S+ to analyze and compare ROC curves. BMC Bioinforma. 12, 77 (2011).

