## Additional files for "Nanopore-based analysis unravels the genetic landscape and phylogenetic relationships of human-infecting *Trichuris incognita* and *Trichuris trichiura* in Côte d’Ivoire, Uganda, Tanzania, and Laos": 270625_Supplementary information.docx

**List of figures**

***Supplementary Fig. 1:*** *Overview of the bioinformatics workflow for genetic diversity study*

***Supplementary Fig. 2:*** *Overview of the quality run characteristics of sequences and reads generated before and after filtering steps with a stringent DADA2 analysis*

***Supplementary Fig. 3:*** *Error plot generated from the DADA2 denoising process run of the sequence analyzed*

***Supplementary Fig. 4:*** *Estimated relative abundance of Trichuris incognita and T. trichiura in mixed infections across countries.*

***Supplementary Appendix 1.*** *DNA extraction protocol for ethanol-preserved Faecal samples*

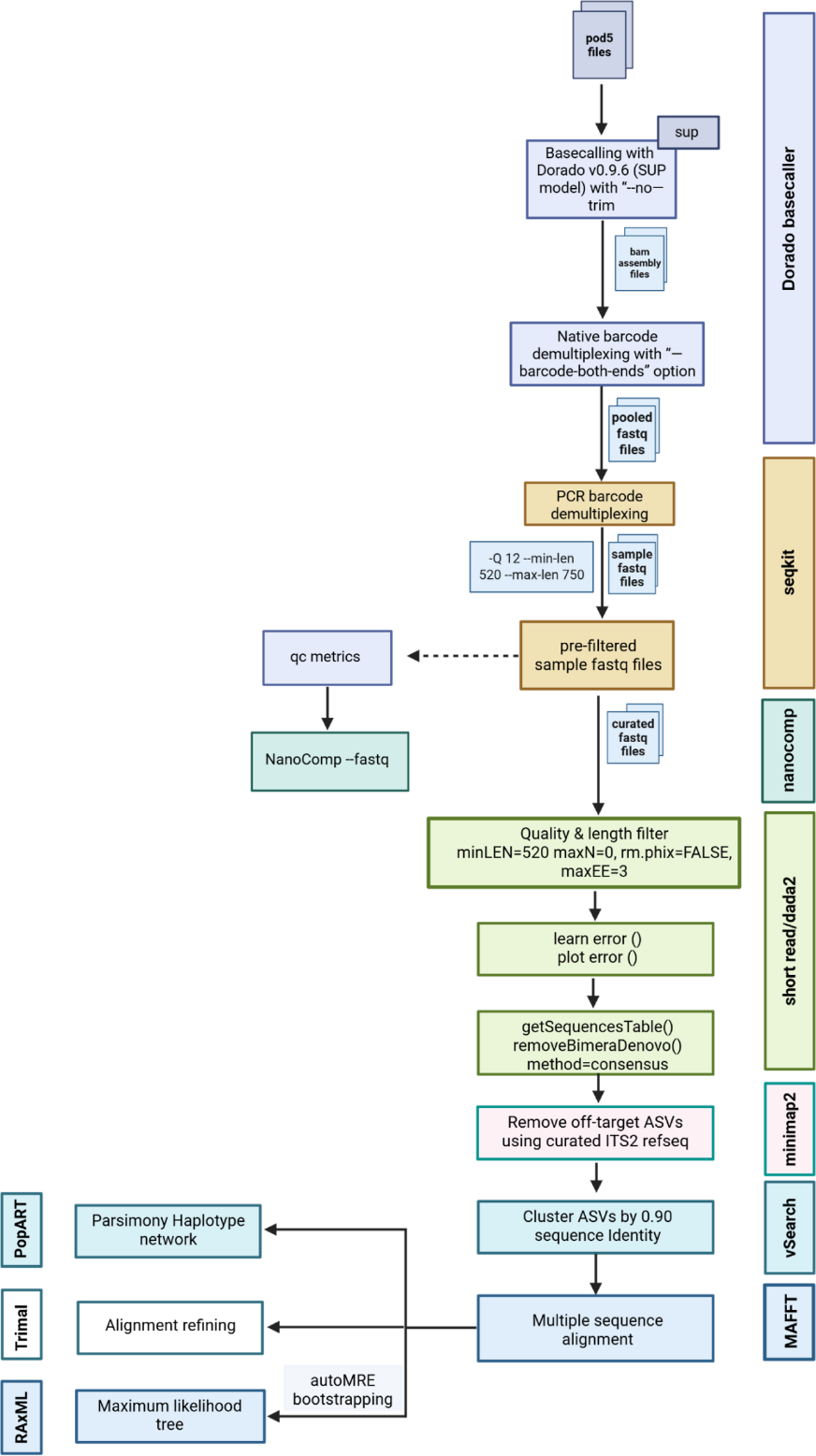

***Supplementary Fig. 1****. Overview of the bioinformatics workflow for genetic diversity study using ONT sequencing data. Softwares used are shown on the right and left ends. Created with BioRender.com.*

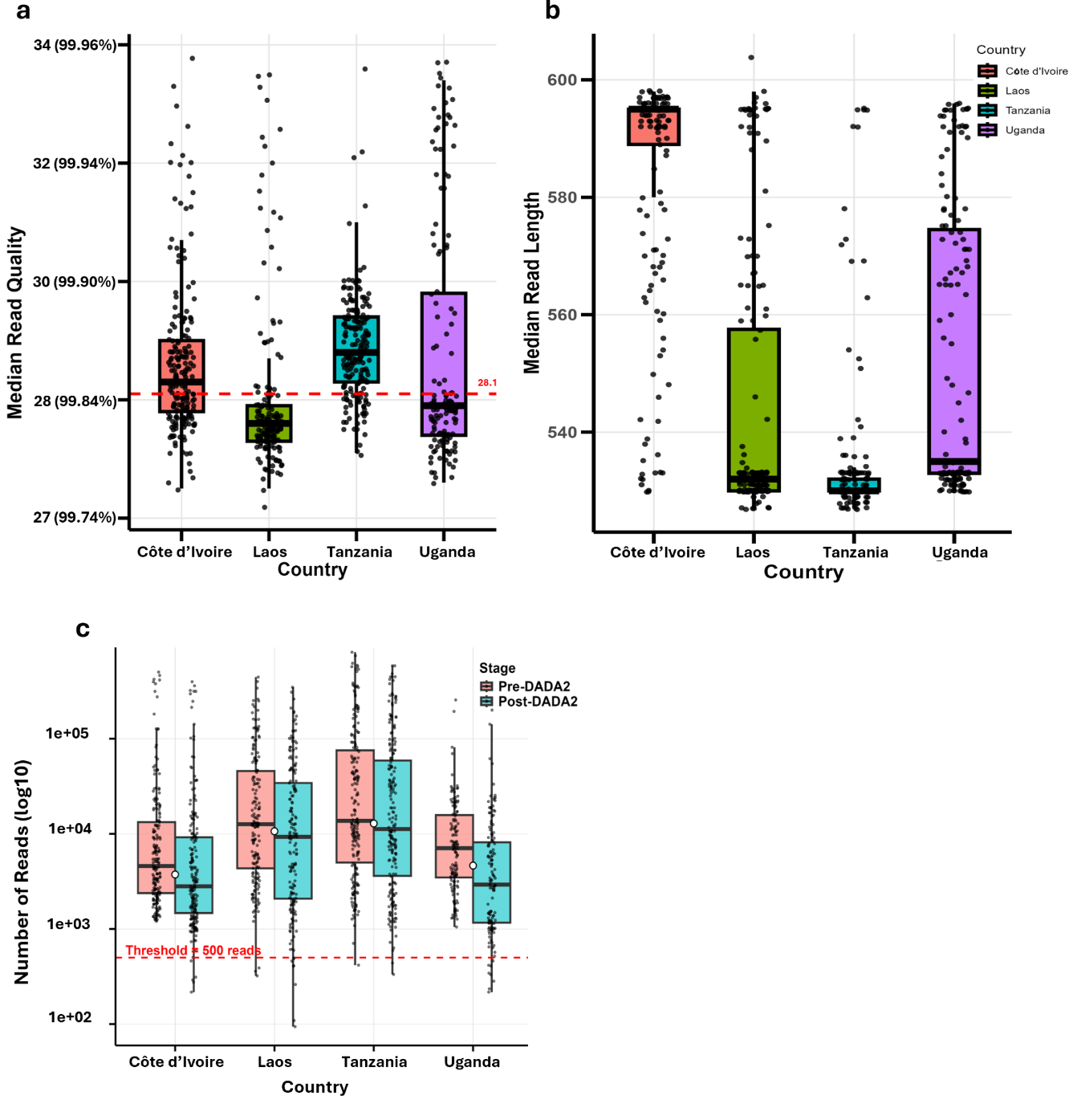

***Supplementary Fig. 2.*** *Overview of the quality run characteristics of sequences and reads generated before and after filtering steps with a stringent DADA2 analysis.****a*** *Median read quality (red line showing the average read quality across all samples sequenced)****. b.*** *Median read length (bp)* ***c.*** *Reads generated before and after DADA2 processing (red line showing the threshold of reads cutoff after a stringent quality filtering approach); Quality (error probability in %), bp = basepairs*

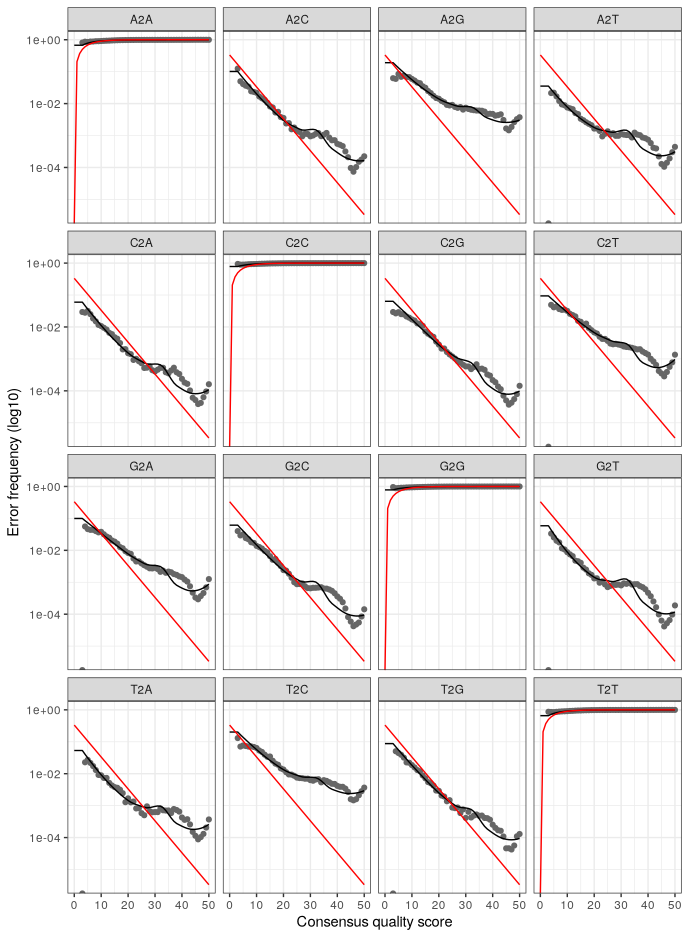

***Supplementary Fig. 3.*** *Error plot generated from DADA2 deniosing process run of sequence analyzed*

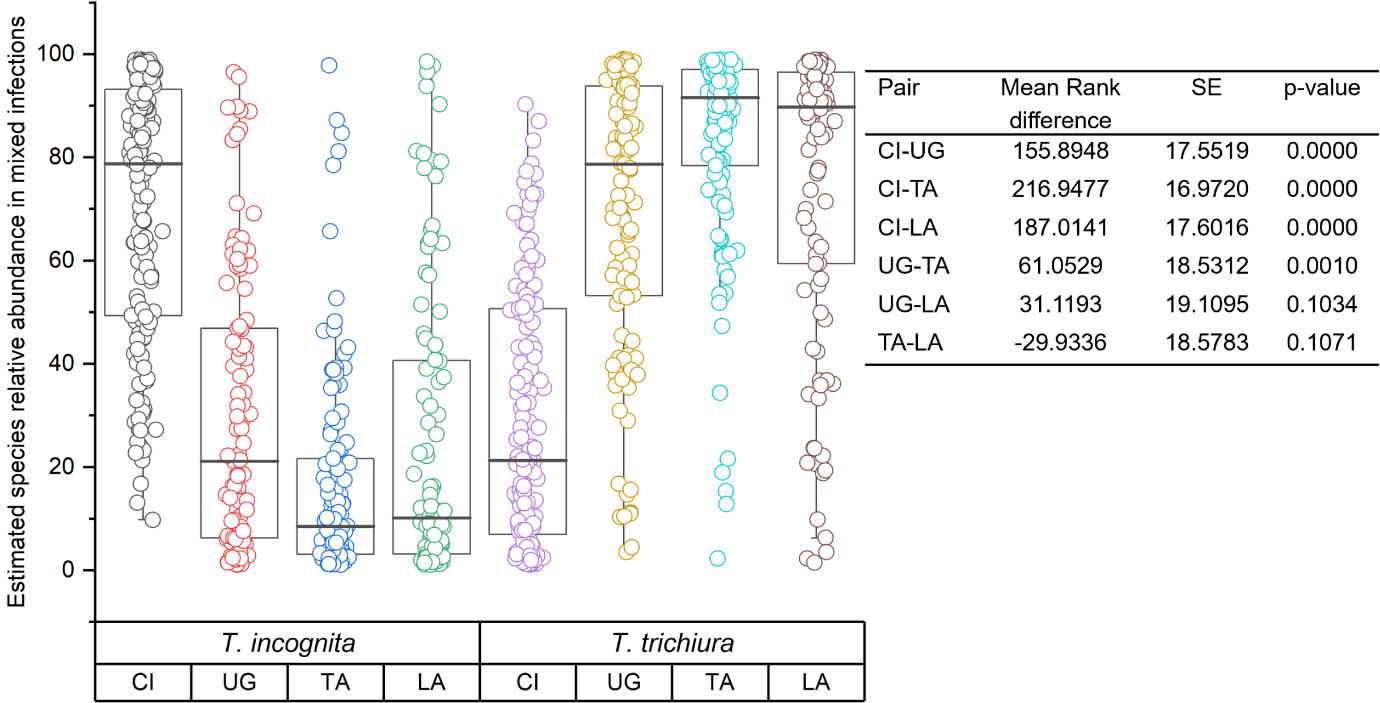

***Supplementary Fig. 4.*** *Estimated relative abundance of Trichuris incognita and T. trichiura in mixed infections across countries. Boxplots show the relative abundance distribution of each species among individuals with mixed infections in Côte d’Ivoire (CI), Uganda (UG), Tanzania (TA), and Laos (LA). Pairwise comparisons of mean ranks between countries for T. incognita are summarized in the embedded table using post-hoc testing (p-values adjusted). Significant differences were observed between CI and all other countries, and between UG and TA.*

***Supplementary Appendix 1.*** *DNA extraction protocol for ethanol-preserved Faecal samples*

1. Mix the ethanol samples with a disposable spatula (homogeneous suspension)

2. Transfer an aliquot of approximately 500 μl of the 96% ethanol-preserved stool sample into an Eppendorf tube (NB: This volume corresponds with the 0.15 – 0.25g of stool recommended by the manufacturer)

3. Centrifuge at 15000 x g for 5 min, and discard the supernatant (ethanol)

4. Wash the pellet by adding 1 ml of phosphate buffer saline (PBS), and mix by vortexing briefly

5. Centrifuge at 15000 x g for 1 min, and discard the supernatant (PBS)

6. Resuspend pellet in 200 μl of PBS

7. Freeze pellet on dry ice for 30 mins

8. Thaw-boil in a preheated thermoblock at 100°C for 10 mins

9. Vortex briefly with the vortex adapter for 3 mins

10. Centrifuge the Powerpro bead tubes at 15000 x g for 1 min

11. Transfer the pellet to Powerpro bead tubes

12. Add 800 µl of Solution CD1 and vortex briefly to mix for 15-20 min

13. Centrifuge at 15000 x g for 2 min

14. Transfer the supernatant to a clean 2 ml Microcentrifuge Tube. Transfer around 800 – 850 μL of supernatant (depending on the nature and volume of the pellet)

15. Add 200 µl of Solution CD2 and vortex for 5 s

16. Centrifuge it at 15000 x g for 1 min. Avoid the pellet, transfer up to 800 μL of supernatant (depending on the volume of your pellet) to a clean 2 mL Microcentrifuge Tube (provided)

18. Add 600 µl of Solution CD3 and vortex for 5 seconds

19. Load 650 µl of the lysate onto an MB Spin Column and centrifuge at 15000 x g for 1 min

20. Discard the flow-through and repeat step 19 to ensure that all of the lysate has passed through the MB Spin Column

21. Place the MB Spin Column into a clean 2 ml Collection Tube and add 500 µl of Solution EA

22. Centrifuge at 15000 x g for 1 min and discard the flow-through

23. Add 500 µl of Solution C5 and centrifuge at 15000 x g for 1 min

24. Place the MB Spin Column into a clean 2 ml Microcentrifuge Tube and centrifuge at 20000 x g for 3 min

25. Place the MB Spin Column into a new 1.5 ml Elution Tube and add 80 µl of Solution C6

26. Centrifuge at 15000 x g for 1 min and discard the MB Spin Column. The DNA is now ready for downstream applications
