## Additional files for "Nanopore-based analysis unravels the genetic landscape and phylogenetic relationships of human-infecting *Trichuris incognita* and *Trichuris trichiura* in Côte d’Ivoire, Uganda, Tanzania, and Laos": Description of Additional Supplementary Files.docx

Files File Name: Supplementary Data 1

Description: Sample metadata and subset metadata for specie-related analysis.

File Name: Supplementary Data 2

Description: Mean Faecal Egg Count (FEC) of the samples. The table shows the Eggs Per Gram (EPG) results of the Kato-Katz analysis for each sample from Côte d’Ivoire, Tanzania, Uganda, and Laos.

File Name: Supplementary Data 3

Description: ASV abundance table, sequences of ASVs generated and taxonomic annotations of samples

File Name: Supplementary Data 4

Description: Haplotype frequencies obtained through the ITS2 sequence analysis of different *Trichuris* populations.

File Name: Supplementary Data 5

Description: List of *Trichuris* haplotypes obtained from humans and their BLAST query. Details on the best match, including accession numbers, percentage of identity, assumed *Trichuris* spp., and the host it was isolated from are provided.

File Name: Supplementary Data 6

Description: Subset metadata for study participants aged 6–18 years.

File Name: Supplementary Data 7

Description: List of primers used for the amplification of *Trichuris* ITS2 rDNA in stool samples. Indicated in bold are the 12 bp long barcodes that were adapted from Dommann et al.^39^ and used for demultiplexing, followed by the ITS2 binding sequence.

File Name: Supplementary Data 8

Description: Sequences of *Trichuris* spp. downloaded from GenBank and used in phylogenetic analysis.
